# Bidirectional transfer of Engrailed homeoprotein across the plasma membrane requires PIP2

**DOI:** 10.1101/2020.01.21.913566

**Authors:** Irène Amblard, Edmond Dupont, Isabel Alves, Julie Miralvès, Isabelle Queguiner, Alain Joliot

## Abstract

1.

Homeoproteins are a class of transcription factors sharing the unexpected property of intercellular trafficking. It confers to homeoproteins a paracrine mode of action, shown to exert a wide range of physiological functions, during development and in the adult. Internalization and secretion, the two steps of intercellular transfer, rely on unconventional mechanisms but the cellular mechanisms at stake still need to be fully characterized. Thanks to the design of new quantitative and sensitive assays dedicated to the study of homeoprotein transfer in cell culture, we demonstrate a core role of the phosphoinositides PIP_2_ together with cholesterol in the translocation of Engrailed2 (EN2) homeoprotein across the plasma membrane. Both secretion and internalization are regulated according to PIP_2_ levels, challenged by drug or enzymatic treatments. In addition, EN2 specifically interacts with PIP_2_ and the reduced affinity of a paracrine deficient mutant of EN2 supports a role of PIP_2_ in homeoprotein physiological function. We propose that the two ways plasma membrane translocation steps accounting for homeoprotein secretion and internalization respectively are parts of a common process.

**Summary Statement:** Deciphering the mechanism of homeoprotein intercellular trafficking

## 2. Introduction

The plasma membrane is a pleiotropic structure that preserves cell homeostasis while allowing the exchange of various molecules, including large biological molecules endowed with signaling properties, mainly proteins. Along this view, the transfer across the plasma membrane constitutes the landmark that characterizes protein internalization and secretion, in opposite directions. In both cases, membrane fusion is the prevalent mechanism, endocytosis for entry and conventional secretion for exit but alternative mechanisms have been proposed.

Conventional protein secretion is initiated by the recognition of a short N-terminal secretion sequence through a co-translational process. In the last decade, unconventional protein secretion (UPS) pathways have been increasingly reported, involving multiple mechanisms (Nickel and Rabouille, 2018; Zhang and Schekman, 2013). Three distinct UPS types are presently proposed (Rabouille, 2017), based on the nature of the secreted substrate (soluble, transmembrane) and of their mode of secretion (plasma membrane translocation, vesicular-based). In particular, UPS Type I refers to soluble proteins that are secreted through direct translocation across the plasma membrane and among them, FGF2 (Schäfer et al., 2004) and HIV Tat (Rayne et al., 2010) are prototypical examples. FGF2 is synthesized as a cytosolic protein but need to be secreted to activate its extracellular receptors. It is devoid of secretion signal but creates pores within the plasma membrane, driving its secretion (Müller et al., 2015). A critical step in FGF2 secretion is its accumulation at the plasma membrane, which requires its specific interaction with PI(4,5)P_2_ (PIP_2_)(Temmerman et al., 2008), a quantitatively minor but essential component of the plasma membrane inner leaflet (Hammond et al., 2012). A similar requirement for PIP_2_ was demonstrated for HIV Tat secretion (Rayne et al., 2010). Although functionally and structurally unrelated, Tat and FGF2 proteins share a high content in basic amino acids (pI of 9.8 and 9.5 respectively).

Protein internalization mostly relies on endocytosis, a vesicular-based pathway initiated by invagination from the plasma membrane (Doherty and McMahon, 2009). Consequently, internalized proteins remain entrapped within the vesicles and never reach the cytosol. Among exceptions, some bacterial toxins are able to translocate through biological membranes and accumulate in the cytosol (Williams and Tsai, 2016). We also learned from the study of Cell Penetrating Peptides that extracellularly loaded protein or polypeptides could have access to the cytosol, although the efficacy and the mechanism(s) involved remain debated (Kauffman et al., 2015).

Intercellular trafficking of homeoproteins constitute a fascinating example of protein transfer across the plasma membrane. These proteins were originally identified thanks to their role in transcriptional regulation and are defined by the nature of their DNA-binding domain, the homeodomain (Gehring et al., 1994). Some years ago, we have observed that homeoproteins present the unexpected property of transfer between cells (Joliot et al., 1998). Indeed, it was recently reported that most if not all homeoproteins are able of transfer (Lee et al., 2019), also emphasizing the pivotal role of the conserved homeodomain in this process (Sagan et al., 2013). Since the original *ex vivo* observations, the physiological significance of homeoprotein intercellular trafficking was revealed (Di Nardo et al., 2018), as it provides to homeoproteins a paracrine mode of action that superimposes on their transcriptional activities. In particular, the paracrine action of Engrailed subclass has been reported in multiple physiological contexts, including drosophila (Layalle et al., 2011), zebrafish (Rampon et al., 2015) and mice (Wizenmann et al., 2009).

Understanding the mechanism of homeoprotein intercellular trafficking remains challenging as both secretion and internalization rely on unconventional mechanisms. Homeoproteins are devoid of secretion signal and predominantly reside in the nucleus. However, homeoproteins such as the chick Engrailed2 homeoprotein (EN2) used as a model homeoprotein, shuttle between the nucleus and the cytosol and around 10 % of the intracellular protein pool is secreted in the soluble fraction of the conditioned medium, thus not incorporated within extracellular vesicles (Maizel et al., 1999). In the cytosol, EN2 associates with the membrane fractions enriched in cholesterol and glycosphingolipids (Joliot et al., 1997). In addition to secretion, homeoproteins are internalized when added in the extracellular medium and accumulates in cytosolic and nuclear compartments, even when endocytosis is inhibited (Joliot et al., 1991). In the present report, we have focused our study on the events taking place at the plasma membrane, the actual barrier that separates the intra and extracellular spaces. We demonstrate that, similarly to FGF2 and Tat, EN2 translocate across the plasma membrane during secretion thanks to its interaction with PIP_2_. Most importantly, reverse translocation during EN2 internalization that lead to its cytosolic accumulation has the same requirement for PIP_2_, suggesting that the plasma membrane translocation events leading to secretion and internalization share crucial steps rather than involving distinct mechanisms.

## 3. Results

### 3.1. PIP2 is required for EN2 secretion

Homeoprotein intercellular trafficking have been best studied with EN2, a polypeptide of 28 kD with a pI of 9.2, it was thus kept for this study. To accurately address the mechanism of protein secretion, we generated stable cell clones in an engineered HeLa line (Tighe et al., 2008) that permits the targeted insertion of the transgene at a invariant site and its inducible expression upon doxycycline addition. Once secreted by EN2 cell clone, EN2 accumulates at the cell surface and can be detected by flow cytometry (Layalle et al., 2011). We verified that EN2 secretion was insensitive to the classical inhibitor of conventional secretion Brefeldin A but became sensitive upon EN2 fusion to an ectopic secretion signal sequence (ssEN2) (Fig. 1A). FGF2 secretion was also insensitive to Brefeldin A to the same extent as EN2 (Fig. S1). Both proteins share a high basicity. Once secreted, they interact with the cell surface through interaction with carbohydrates and both proteins were efficiently removed by heparin wash treatment (Fig. 1B).

**Figure 1:**
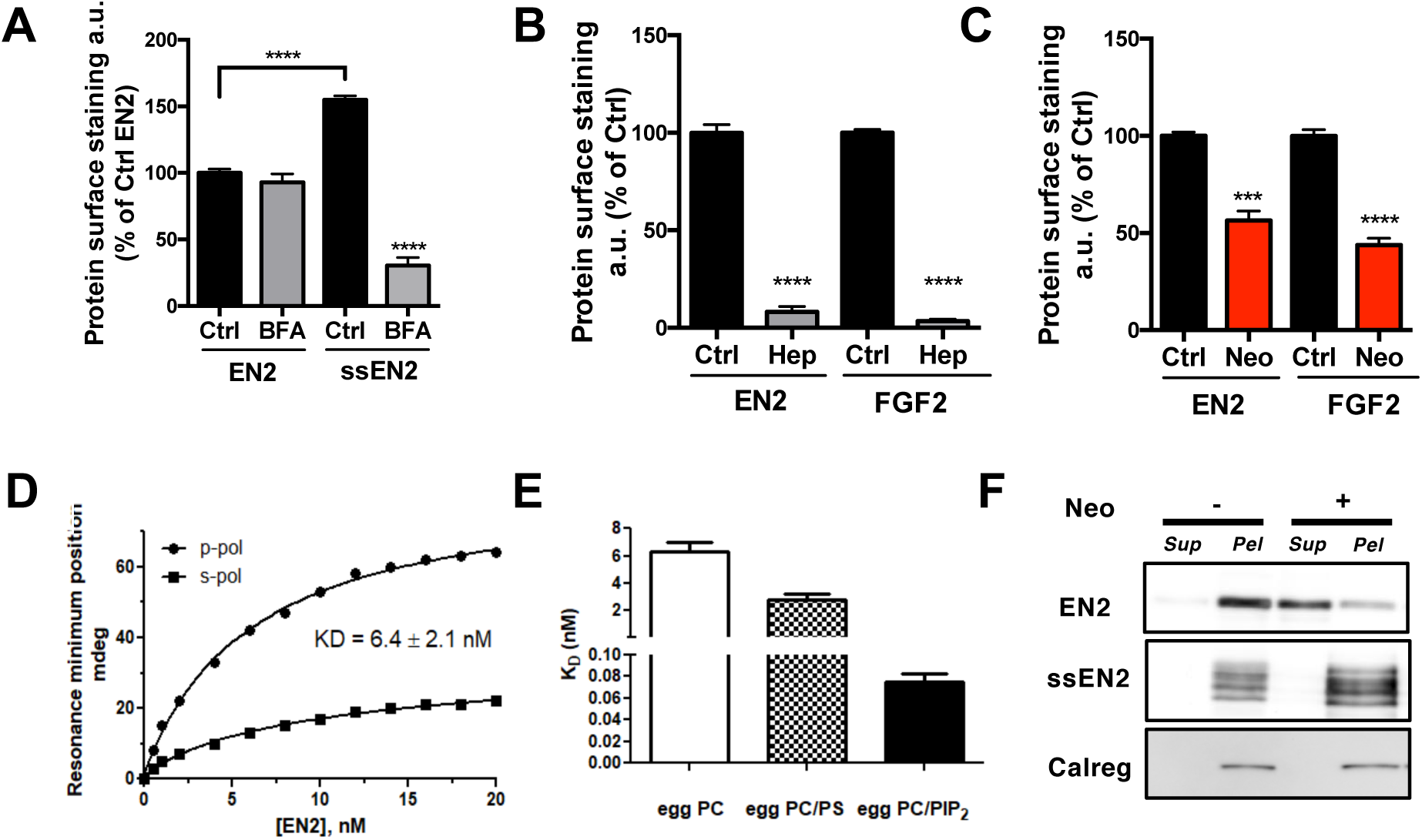
PIP_2_ is required for EN2 unconventional secretion and enhances lipid membrane affinity. (A) Cell surface accumulation of EN2 or ssEN2 treated or not with Brefeldin A (BFA) for 2h. (B) Cell surface removal of EN2 or FGF2 following heparin treatment (Hep) for 5 min. (C) Cell surface accumulation of EN2 or FGF2 in the presence or not of 10 mM neomycin (Neo) for 18h. (D,E) Interaction of EN2 with artificial planar lipid bilayers composed of egg PC, egg PC/DOPS (3/1 mol/mol) and egg PC/PIP_2_ (9/1 mol/mol) measured by PWR. Each experiment was repeated 3 times. (D) Changes in the minimum resonance position for both p (●) and s (▪) polarized time as a function of concentration of EN2 from which KD values have been determined. (E) presents the KD values obtained from 3 independent experiments. (F) Western blot analysis of the release of EN2, ssEN2 and calregulin from the membrane fraction treated or not with 10 mM neomycin. Once separated by centrifugation, equivalent fractions of supernatant (Sup) and Pellet (Pel) fractions were analyzed.

Since it was demonstrated that FGF2 interaction with the negatively charged Phosphatidylinositol-4,5-bisphosphate (PIP_2_) is required for its secretion (Temmerman et al., 2008), we asked whether the PIP_2_-interacting drug neomycin known to inhibit FGF2 secretion also impaired EN2 secretion. Overnight neomycin treatment inhibited EN2 and FGF2 secretion to similar extent (Fig. 1C), indicating the involvement of PIP_2_ in EN2 secretion. We then evaluated whether EN2 directly interacts with PIP_2_ as demonstrated for HIV Tat (Rayne et al., 2010) and FGF2 (Temmerman et al., 2008). The interaction of EN2 recombinant protein with lipid bilayers of controlled composition was analyzed by Plasmon-Waveguide Resonance (PWR) (Harté et al., 2014), a spectroscopic technique allowing to directly measure the interaction and affinity of molecules with membrane lipids. Significant interaction was observed (Fig. 1D) with pure phosphatidylcholine (PC) bilayer (KD:6.3 nM) but incorporation of PIP_2_ (10%) dramatically lowered the KD to 75 pM, compared to the mild effect elicited by addition of phosphatidylserine (PS) (KD:2.8 nM) (Fig. 1E). As expected, the specific interaction between FGF2 and PIP_2_ was confirmed using the same experimental set-up, although with slightly lower affinity compared to EN2 (Fig. S1).

We have previously shown that a pool of EN2 associates with the membrane fraction, both *in vivo* and *ex vivo* (Joliot et al., 1997). We thus asked whether this association depends on PIP_2_ interaction. The membrane fraction of HeLa EN2 or ssEN2 cells clones was incubated with neomycin and the protein released in the medium was separated from membranes by ultracentrifugation. More than 50 % of the EN2 protein was retrieved in the supernatant following neomycin treatment (Fig. 1F), highlighting the involvement of PIP_2_ in this association. Neither ssEN2 nor the endoplasmic reticulum marker calregulin distributions were affected by this treatment, attesting vesicle integrity and consequently, that the EN2 pool sensitive to neomycin was not entrapped within vesicles.

### 3.2. A new assay for the analysis of unconventional secretion

As PIP_2_ mostly reside in the inner leaflet of the plasma membrane, we decided to focus on this compartment. Flow cytometry, although quantitative at a single cell level, is poorly informative on the mechanism of secretion. To address this question, we took advantage of the Rush strategy developed by Boncompain et al. (Boncompain and Perez, 2012), an inducible secretion assay. A secreted substrate fused to a streptavidin binding peptide (SBP) is sequestered by the co-expression of a “hook” protein fused to core streptavidin (Strep). Biotin addition induces the synchronized release of the substrate from the hook and its eventual secretion. Importantly, the subcellular localization of the hook determines the site of release along the secretion pathway of the substrate upon biotin addition. We chose a hook that localizes at the inner side of the plasma membrane, by Strep fusion either with the acylation sequence of the LCK protein or at the cytosolic side of a transmembrane protein fragment (TM1). Flow cytometry was not sensitive enough to monitor the secretion burst induced by biotin addition (not shown). We thus implemented our assay with a highly sensitive detection tool (HiBit assay, Promega), based on light production upon spontaneous complementation of two Nanoluciferase fragments.

To monitor protein secretion, the small 11 aa fragment (HiBiT) was fused to the secreted substrate and the large one (LgBiT) was expressed at the extracellular side plasma membrane upon fusion at the N-terminus of the TM1Strep trans-membrane protein. By combining the two strategies (Fig. 2A), secretion of the pool of EN2 sequestered at the cytosolic side of the plasma membrane was measured by the specific increase in light production induced by biotin addition compared to control conditions (without biotin). To minimize non-specific release, stable cell clones were generated for all secreted substrates in the FlpIn-TREX HeLa cell line. LCK-StrepCherry (hook) was transiently expressed in SBP-EN2-Hibit (named EN2* for convenience) cell clone. By immunofluorescence we verified on permeabilized cells the retention of EN2* at the plasma membrane in transfected cells (white arrows) and its release by biotin addition (Fig. 2B). Concomitantly, EN2* cell surface staining (visualized in non-permeabilized cells) was increased by biotin treatment (Fig. 2C). A similar behavior was observed with the FGF2* cell clone (Fig S2).

**Figure 2:**
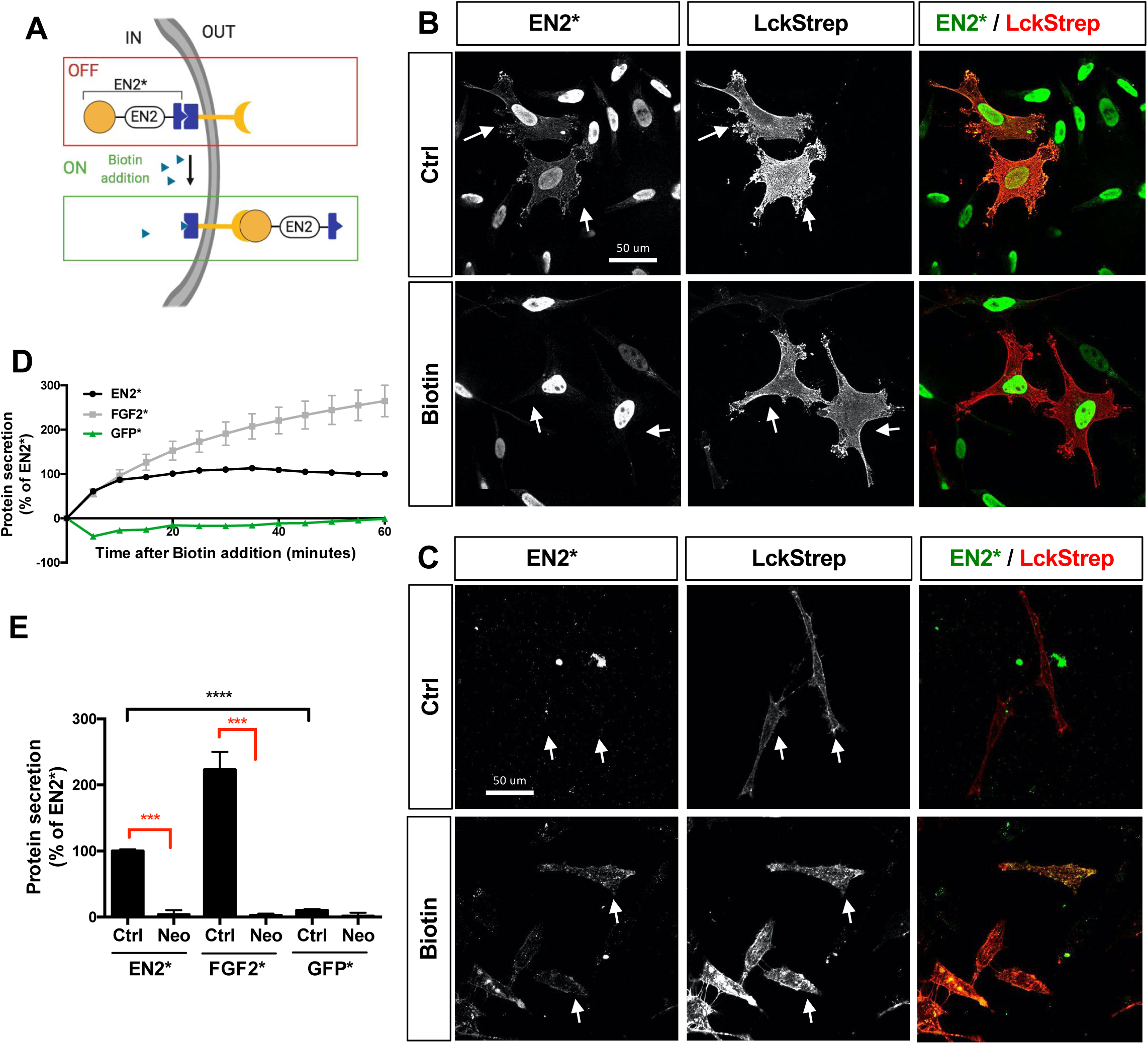
Assessing EN2 unconventional secretion. (A) Principle of the TransRush assay for measuring EN2 secretion based on the combination of RUSH (SBP/streptavidin) and HiBit (Hibit/LgBit) assays. (B,C) EN2* cell line was transfected with a LCKStrepCherry construct and treated or not with biotin (60 min, 100 µM). (B) Intracellular or (C) extracellular EN2* was detected by immunofluorescence using permeabilized or unpermeabilized conditions respectively. (D) Kinetics of biotin-induced secretion of EN2* (black), FGF2* (grey) and GFP* (green) proteins. (E) Quantification of EN2*, FGF2* and GFP* protein secretion in cells treated (Neo) or not (Ctrl) with neomycin (10 mM) for 1 hour.

The secretion of three different proteins was then analyzed by luminescence measurements, GFP* and FGF2* being used respectively as negative and positive controls. The hook (LCK-StrepCherry and the sensor/hook (LgBitTM1Strep) were co-expressed in each clone and following biotin addition together with the luciferase substrate furimazin, the kinetics of secretion was monitored for one hour. Cells were then lysed and the intracellular content for each protein was measured by luminescence for normalization. Subtracting biotin-negative from biotin-positive measurement allowed us to remove the contribution of non-sequestered proteins (e.g. expressed in non-transfected cells). Secretion-induced luminescence started right after biotin addition with FGF2* and EN2*clones, supporting a direct translocation of the intracellular hooked proteins. In the same conditions, no GFP* secretion could be observed (Fig. 2D), thus validating our secretion assay which was named TransRush assay (Rush targeting translocation). We also noticed a higher efficiency of FGF2* secretion compared to EN2* one. An additional benefit of our assay is to allow the measurement of secretion on a short time scale, more suitable to analyze the impact of pharmacological treatments while limiting indirect or toxic effects. We first tested the effect of short-time neomycin treatment (added 30 min before biotin addition) in the TransRush assay. One hour after biotin addition, both EN2* and FGF2* secretions were drastically reduced by neomycin treatment to similar extents, further supporting a direct action of the drug at the site of the plasma membrane (Fig. 2E).

### 3.3. PIP2 levels modulate EN2 secretion

We next asked whether EN2 secretion was sensitive to PIP_2_ levels, using an enzymatic strategy. OCRL is a 5-phosphatase that dephosphorylates PIP_2_ into PI(4)P and its ectopic expression drastically reduced PIP_2_ levels, as revealed with anti-PIP_2_ antibodies (Fig. 3A). The CherryOCRL coding sequence was inserted in a bi-expressor plasmid together with the LgBiTM1Strep (sensor/hook) sequence to maximize their co-expression. Cherry-OCRL protein, but not Cherry co-expression, reduced EN2* and FGF2* secretion after biotin addition (Fig. 3B). Conversely, we expressed the PI4P5K1*α* enzyme to increase PIP_2_ levels at the plasma membrane. As shown in Fig 3C, cells expressing Cherry-PI4P5K showed a moderate but consistent increase of PIP_2_ levels and a concomitant increase of of both EN2* and FGF2* secretions (Fig 3D). Taken together, these results demonstrate the pivotal role of PIP_2_ levels in EN2 unconventional secretion.

**Figure 3:**
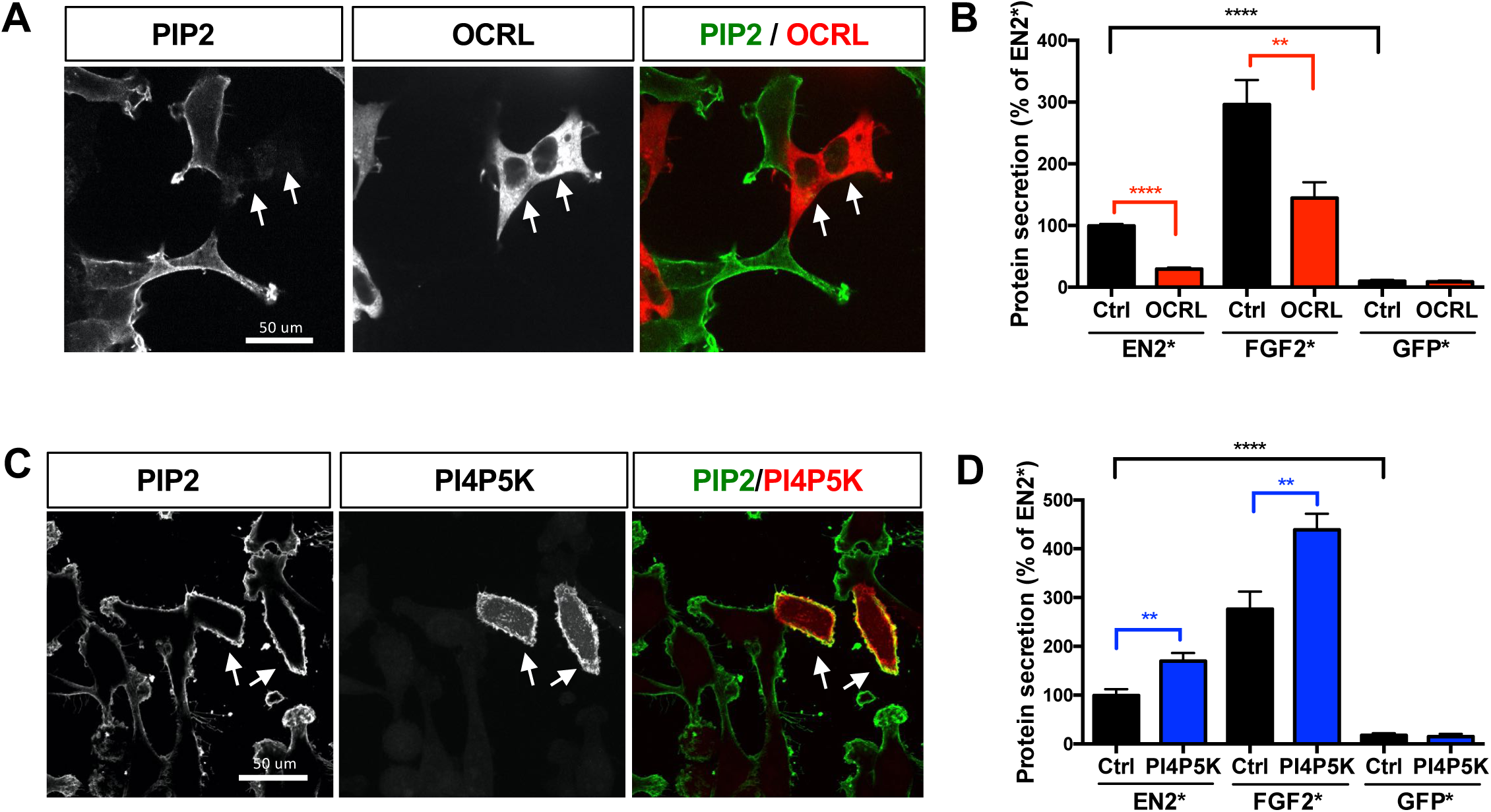
PIP_2_ directly modulates the kinetics of EN2 and FGF2 secretion. (A) Immunostaining of PIP_2_ (green) in cells expressing CherryOCRL (red). (B) Quantification of biotin-induced EN2*, FGF2* and GFP* protein secretion in cells expressing Cherry (Ctrl) or CherryOCRL (OCRL), measured 1 hour after biotin addition. (C) Immunostaining of PIP2 (green) in cells expressing Cherry-PI4P5K (red). (D) Quantification of biotin-induced EN2*, FGF2* and GFP* protein secretion in cells expressing Cherry (Ctrl) or Cherry-PI4P5K (PI4P5K), measured 1 hour after biotin addition.

### 3.4. PIP2 modulates Engrailed2 internalization

Homeoproteins are not only secreted but are also internalized and reach cytosolic and nuclear compartments. After a one-hour incubation in the culture medium, FITC-labelled recombinant EN2 (F-EN2) was mainly detected at the cell surface (Fig 4A, Ctrl). The specific intracellular staining was revealed after quenching all the extracellular fluorescence with trypan blue (Vranic et al., 2013). Both punctuate and diffuse distributions were observed (Fig 4A, +TB), the latter highlighting the cytosolic and nuclear accumulation of the protein. As PIP_2_ are required for EN2 secretion, we asked whether they would also be involved in the reverse process of internalization. In cells treated with neomycin, F-EN2 intracellular accumulation was drastically reduced (Fig. 4B). Importantly, the extracellular association of F-EN2 with the plasma membrane, detected before trypan blue addition, was not affected by neomycin treatment (Fig S3), supporting an intracellular action of the drug. Next, we tested the effect of PIP_2_-modulating enzymatic treatments. Hela cells were transfected with Cherry fusion constructs and F-EN2 internalization was compared between transfected (Cherry positive) and non-transfected cells. Cherry-OCRL expression significantly impaired F-EN2 internalization (Fig. 4C). Using the same protocol, Cherry-PI5P4K expression increased F-EN2 internalization (Fig 4D).

**Figure 4:**
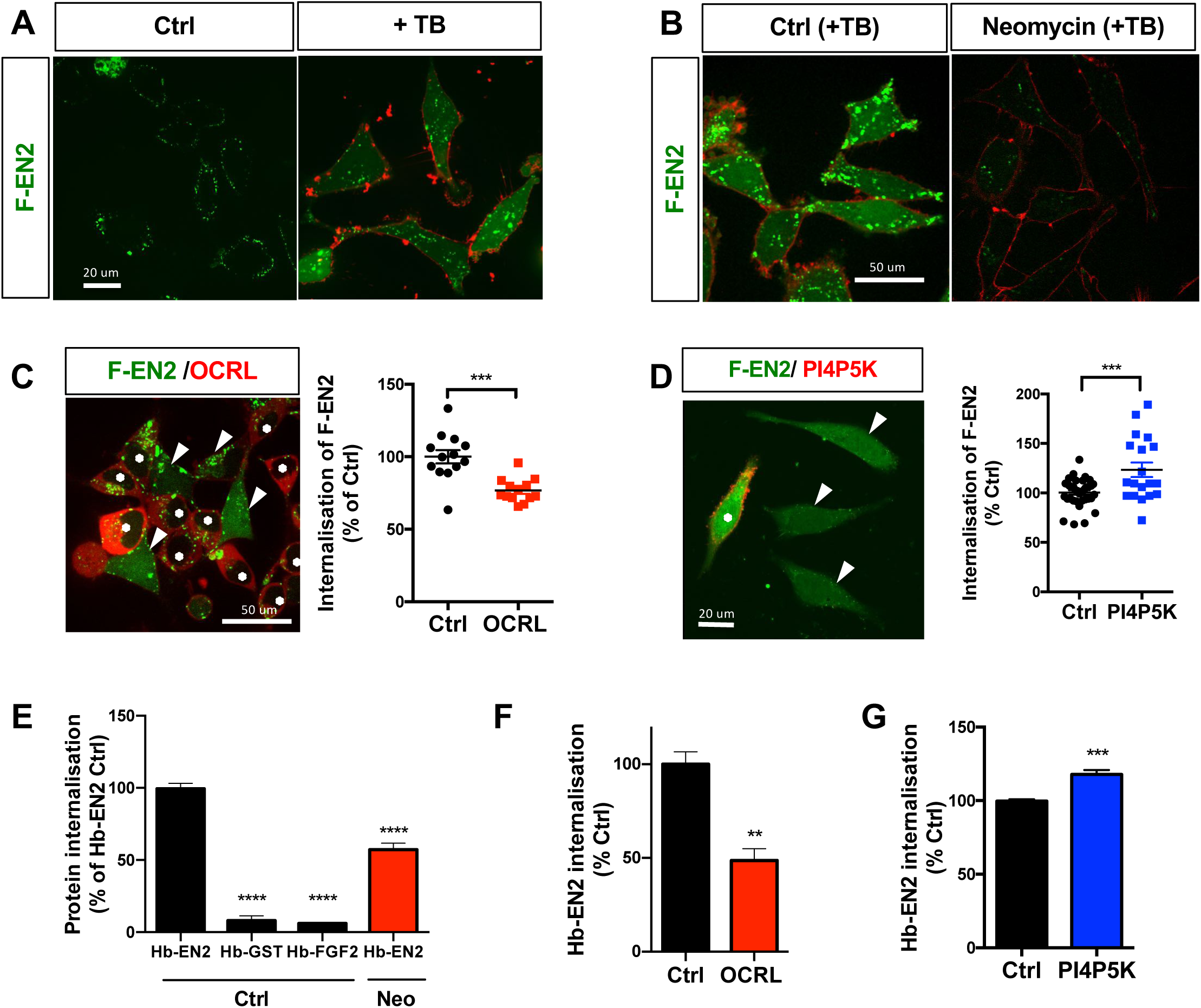
PIP_2_ modulate EN2 internalization. (A) F-EN2 (2 µM) staining in live cells following 1 hour incubation before (Ctrl) or after (+TB) trypan blue addition (0.1% final concentration). Intrinsic trypan blue fluorescence is visualized (red). (B) Intracellular F-EN2 (2uM) staining in live cells after 1 hour incubation in the presence (Neomycin) or absence (Ctrl) of 10 mM after TB addition. (C,D) Intracellular F-EN2 (2uM) staining following 1 hour incubation in live cells expressing (C) Cherry-OCRL or (D) Cherry-PI4P5K after TB addition. Diffuse staining intensity is quantified in cells expressing (*) or not (arrowhead) the respective enzymes (E-G) Luminescence-based quantification of Hb-EN2, Hb-FGF2 and Hb-GST internalization following one hour incubation with cells, (E) treated or not with 10 mM neomycin (Neo), or expressing (F) CherryOCRL or (G) Cherry PI4P5K. Cherry expressing cells (Ctrl) were used as a control in (F) and (G).

Because a distinctive feature of plasma membrane translocation-driven processes is the cytosolic accumulation of the internalized protein, we have adapted the HiBiT assay to selectively target this mode of internalization. We first added a HiBiT tag to the recombinant EN2 protein (Hb-EN2) and secondly generated a reporter cell clone that expressed the LgBiT protein in the cytosol. Light production only occurs if extracellularly added Hb-EN2 reaches the cytosol, which can be measured in live cells thanks to the quick diffusion of the luciferase substrate into live cells (Song et al., 2013). One hour after its addition in the extracellular medium, Hb-EN2 efficiently stimulated light production together with the cytosolic reporter (Fig. 4E). As expected, neither Hb-GST nor Hb-FGF2 could activate the reporter. Both neomycin treatment (Fig. 4E) and OCRL ectopic expression (Fig. 4F) lowered the cytosolic accumulation of Hb-EN2, whereas PI4P5K ectopic expression had an opposite enhancing effect (Fig 4G).

### 3.5. Selective role of raft associated PIP2 in EN2 trafficking

In a previous study, we have shown that EN2 selectively associates with cholesterol-enriched membranes (Joliot et al., 1997). Using the PWR assay, we demonstrated that cholesterol incorporation in the membrane increased 10 fold EN2 affinity for a PIP_2_ containing lipid bilayer (Fig 5A). We first asked whether cholesterol levels directly modulate EN2 intercellular trafficking. Extracting cholesterol from the plasma membrane by methyl-ß-Cyclodextrin (MCB) treatment reduced F-EN2 internalization (Fig. 5B) and EN2* secretion (Fig. 5C), indicating that similarly to PIP_2_, cholesterol contribute to EN2 bi-directional plasma membrane trafficking.

**Figure 5:**
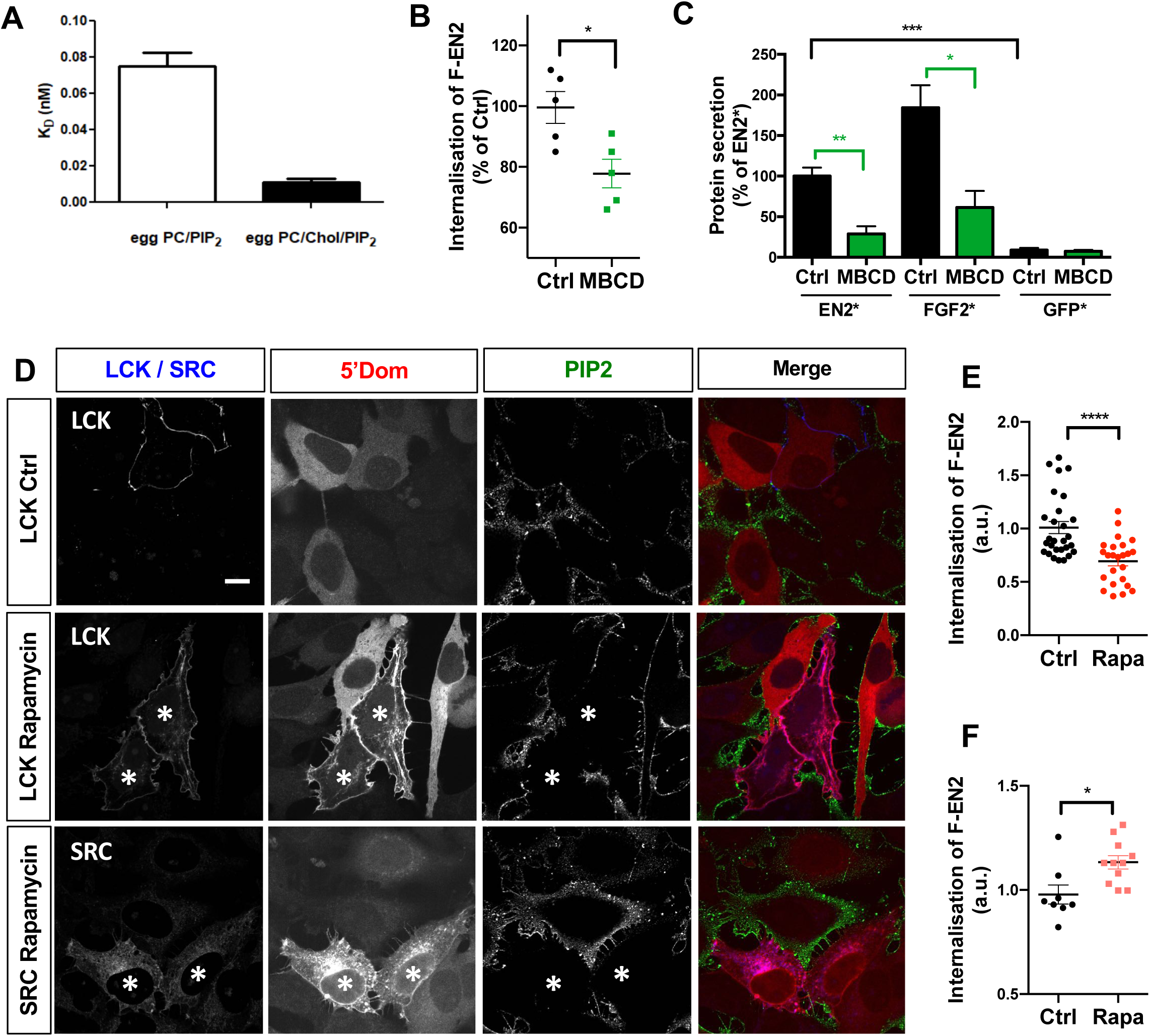
Role of Cholesterol and raft-associated PIP_2_. Interaction and affinity of EN2 with planar artificial lipid bilayer composed of egg PC/chol/PIP_2_ (7/2/1 mol/mol/mol) determined by PWR, the condition with egg PC/PIP_2_ (9/1 mol/mol) is identical to Fig. 1E. Each experiment was repeated 3 times. (B) Intracellular F-EN2 (2 µM) staining in live cells following 1 hour incubation in the presence or not of 10mM cyclodextrin (MBCD). (C) Quantification of biotin-induced EN2*, FGF2* and GFP* protein secretion in cells pre-treated or not with 10 mM MBCD for 30 minutes. (D) Immunofluorescence detection of PIP_2_ (green) in cells co-expressing FKBP-Cherry5Dom (red) and CFP-FRB fused to SRC or LCK anchor in cells treated (Rapamycin) or not (ctrl) with rapamycin 100nM for 30 min. (E,F) Quantification of F-EN2 intracellular staining in cells co-expressing FKBP-Cherry5Dom and CFP-FRB fused to LCK (E) or SRC (F) anchor and treated or not with 100 nM rapamycin for 30 min(Rapa).

PIP_2_ are known to associate with rafts, the cholesterol enriched membrane domains (Pike and Casey, 1996). We verified that PIP_2_ co-purified with the raft fraction isolated by density gradient separation of detergent-treated membranes (Fig S4). We asked whether this PIP_2_ pool was specifically involved in EN2 trafficking. To selectively deplete raft-associated PIP_2_, we used an assay developed by Varnai et al.(Varnai et al., 2006), which consists in the selective targeting of the cytosolic phosphatase domain of the TypeIV-5Phosphatase (5Dom). LCK or SRC acylation motifs that segregate towards toward raft or non-raft domains. respectively (Fig S4), were used to drive 5Dom relocalization using the FRB/FKBP rapamycin-induced dimerization assay. Plasma membrane targeting of 5Dom by both LCK and SRC anchors lowered PIP_2_ levels (Fig. 5D) but only raft-targeted 5Dom significantly impaired F-EN2 internalization (Fig. 5E). By contrast, non-raft PIP_2_ depletion induced a small increase in EN2 internalization (Fig. 5F). Such depletion was previously shown to increase the raft-associated PIP_2_ pool (Johnson et al., 2008). We concluded that cholesterol and the raft associated PIP_2_ pool cooperate to regulate EN2 intercellular trafficking.

### 3.6. Competing for PIP_2_ interaction in live cells

HIV Tat, whose secretion also depends on PIP_2_, efficiently delocalizes PHPLC from the plasma membrane (Rayne et al., 2010). We thus decided to follow the behavior of the PIP_2_-interacting probe GFP-PHPLC∂ during EN2 trafficking. EN2 uptake quickly induced a delocalization of GFP-PHPLC∂ probe toward the cytosol (Fig. 6A), but not that of a LCK fusion protein used as control (Fig. S5). Redistribution started only few minutes after EN2 addition and increased with time (Fig. 6B) We then asked if EN2 secretion also promotes PHPLC∂ delocalization. GFP-PHPLC∂ was expressed in the TransRush assay. Promoting EN2* or FGF2*secretion by biotin addition induced a delocalization of GFP-PHPLC∂ towards the cytosol similarly to EN2 internalization (Fig. 6C and 6D). These two results further confirm that PIP_2_ is an essential relay in EN2 translocation across the plasma membrane in either direction. Using the same experimental protocol, FGF2* secretion also delocalized GFP-PHPLC∂ upon biotin addition (Fig. 6D) but had little effect when applied in the extracellular medium, in agreement with its inability to enter cell by direct translocation (Fig. 4E).

**Figure 6:**
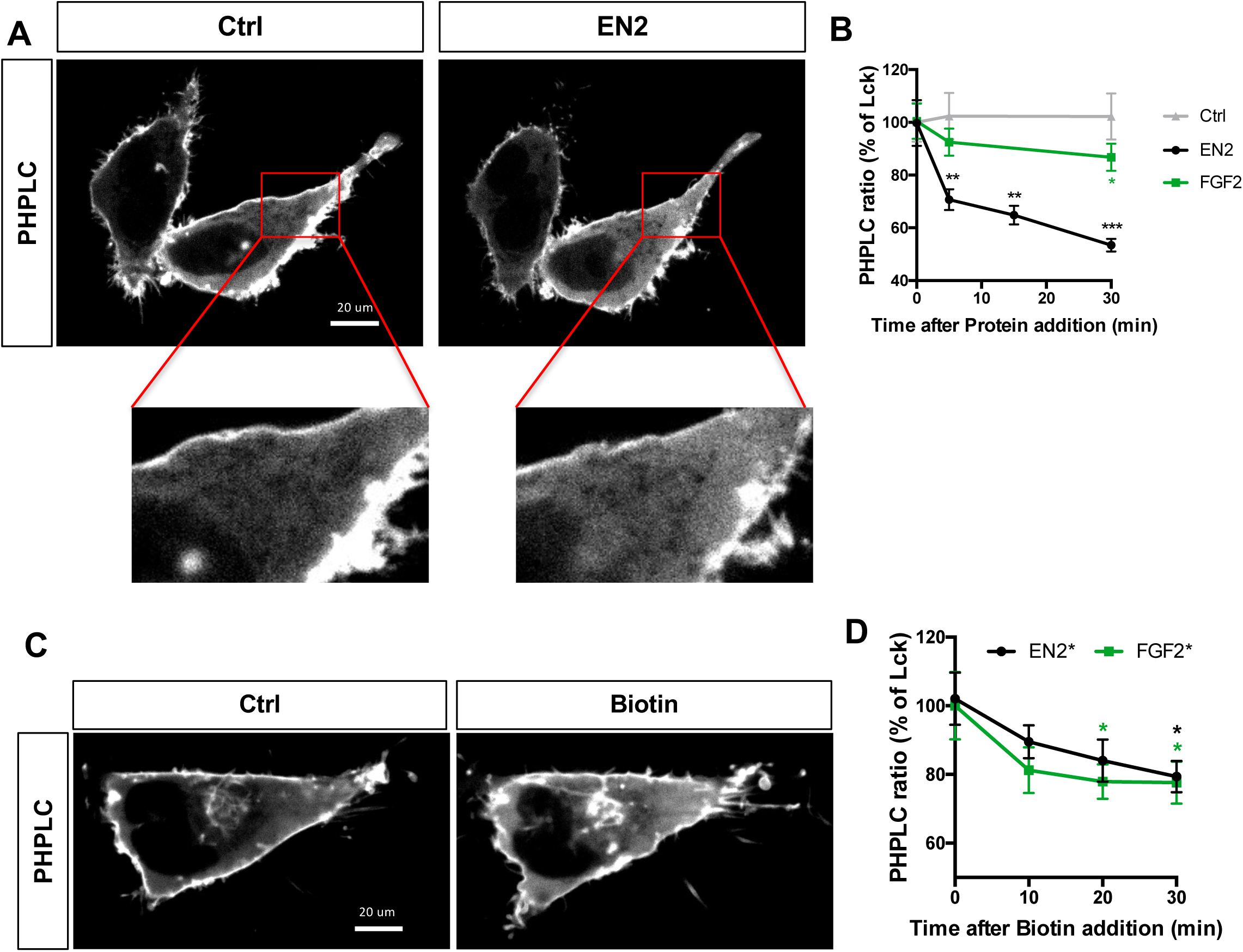
EN2 trafficking induces PHPLC delocalization. (A) PHPLC-GFP distribution in cells before (ctrl) and after (EN2) incubation of extracellular 2 µM EN2 protein for 30 min. (B) Kinetics of PHPLC delocalization following addition of 2 µM EN2 or FGF2 proteins (C) PHPLC-GFP distribution in EN2* cell clone before (ctrl) and after induction of EN2* secretion with biotin (Biotin) for 30 min (D) Kinetics of PHPLC delocalization following induction of protein secretion in EN2* or FGF2* cell clone.

### 3.7. A paracrine-deficient mutant of Engrailed has a reduced affinity for PIP2

Intercellular transfer confers to homeoproteins a paracrine mode of action with strong physiological outcomes. In zebrafish, we have shown recently that the paracrine activity of EN2 protein ectopically expressed in embryo induces eye defects (Rampon et al., 2015). We have characterized a paracrine-deficient mutant of EN2 (EN2_ww>KK_), which was both unable to induce such phenotype and strongly impaired in its intercellular trafficking. We first confirmed using the TransRush assay the altered behavior of the EN2_ww>KK_ mutant. Both internalization (Fig 7A) and secretion (Fig.7B) were significantly impaired. Additionally, the EN2_ww>KK_ mutant was unable to delocalize PHPLC in either situation (Fig 7C, D). We thus asked whether the interaction of EN2_ww>KK_ with PIP_2_ was concomitantly affected by the mutation. Compared to the wild type EN2 protein, the affinity of EN2_ww>KK_ mutant for a pure PC-containing membrane (KD:20 nM) was not significantly reduced but a high impact was observed with membrane containing PIP_2_ with an almost 30 fold decrease in affinity (KD:2.5 nM) (Fig 7E). These results further support the pivotal role of PIP2 interaction not only in EN2 trafficking but also in the paracrine activity of the protein.

**Figure 7:**
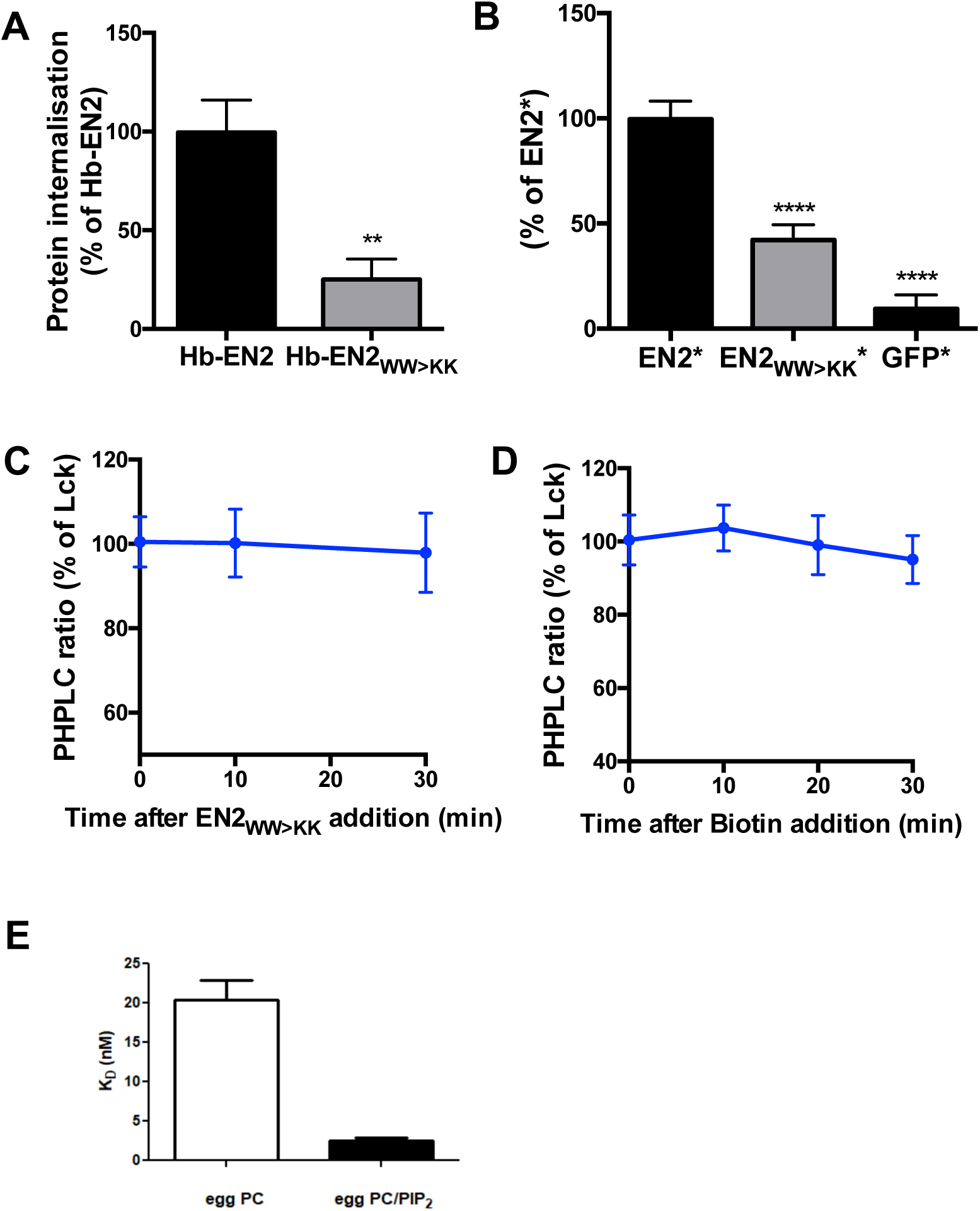
The impaired ability to transfer of EN2_WW>KK_ mutant correlates with its reduced affinity for PIP_2_. Analysis of EN2_WW>KK_ mutant in (A) internalization and (B) secretion assays. Quantification of GFP-PHPLC delocalization following (C) extracellular addition of 2 µM EN2 _WW>KK_ and (D) upon stimulation of EN2 _WW>KK_ * secretion by biotin addition (E) Interaction and affinity of EN2_WW>KK_ with artificial planar lipid bilayers composed of egg PC and egg PC/PIP2 (9/1 mol/mol) measured by PWR

## 4. Discussion

In the present study, we have investigated the molecular and cellular mechanisms accounting for the intercellular transfer of homeoproteins, concentrating on the crossing of plasma membrane. The two steps of the transfer, secretion and internalization, have been analyzed separately in this study. Combining Rush and HibiT assays in the TransRush assay, we demonstrate that a pool of EN2 that has reached the inner side of the plasma membrane is secreted by direct translocation towards the extracellular medium. We further demonstrate the specific interaction of EN2 with PIP_2_ and its pivotal role in secretion. First, the extent of secretion strictly correlates with PIP2 levels, modulated either by enzymatic treatments or drugs. Secondly, EN2 compete for PIP_2_ binding with PHPLC∂ when secreted. EN2 is a new example of PIP_2_-dependent secreted substrate along with FGF2 and HIV Tat, both synthesized in the cytosol and highly basic. EN2 secretion would thus correspond to a UPS Type I pathway, although the molecular mechanism of the translocation remains to be determined. Using a dedicated assay that selectively targets the cytosolic accumulation of the internalized protein, we show here that this mode of internalization shares with secretion a very same requirement for PIP_2_, strongly suggesting that these two events are part of a same process, both involving the translocation of the protein across the plasma membrane but in two opposite directions. EN2 secretion and internalization might thus result from a dynamic equilibrium at either side of the plasma membrane with the local micro-environment.

In a more general way, translocation of soluble proteins across the plasma membrane raises the question of their adaptation to two radically different environments, the hydrophilic cytosol and the hydrophobic core of the membrane. At the plasma membrane, FGF2 and Tat proteins assemble into oligomers which enable the formation of pores (Müller et al., 2015; Zeitler et al., 2015). We have no evidence of multimer formation in the case of EN2, which possess only one cysteine. However, in the presence of negatively charged lipid bicelles, the conformation of EN2 protein analyzed by NRM is shifted, promoting the exposure of the hydrophobic core of the homeodomain (Carlier et al., 2013). The interplay between electrostatic and hydrophobic (supported by the reduced affinity of the EN2_ww>KK_ mutant) interactions with PIP_2_ might act as a switch for EN2 to shuttle between the two environments. The highly negatively charged glycosaminoglycans that concentrate at the cell surface might play a similar role on the extracellular side of the plasma membrane. Indeed, they are required for FGF2 secretion (Zehe et al., 2006) and dictates the specificity of Otx2 homeoprotein internalization *in vivo* (Beurdeley et al., 2012).

The paracrine mode of action of homeoprotein is a direct consequence of their trafficking properties. First demonstrated with selected homeoproteins(Di Nardo et al., 2018), a recent study that now shows that intercellular trafficking is common to most if not all homeoproteins (Lee et al., 2019), suggests that paracrine functions might be a shared property within this protein family. Whether PIP_2_ might act as a generic regulator of homeoprotein trafficking is an interesting possibility, also considering that PIP_2_ levels are the target of multiple signalization pathways within the cell.

## 5. Methods

### 5.1. Cell culture

Cell culture experiments were performed on HeLa cells grown in DMEM supplemented with 10% fetal bovine serum. Transient transfections were performed with Lipofectamine 2000 (Life Technologies) according to the manufacturer’s instructions. Cells were cultured for an additional 24 h before being processed for analysis. Stable clones were generated for each protein used in the secretion assays. All cell clones were produced in a modified HeLa cell line (HeLa FlpIn-TREX) kindly provided by S. Taylor (Tighe et al., 2008) according to the instructions of the FlpIn-TREX Core Kit (Invitrogen) and are listed in Table S3. Briefly, the cell line allows the targeted insertion of the transgene by co-expression of the FlpIn recombinase as well as its inducible expression by doxycycline when using appropriate promoter thanks to the constitutive expression of the tetracycline repressor.

### 5.2. Chemicals and Pharmacological treatments

All drugs and Chemicals were purchased at Sigma unless specified. To inhibit conventional protein secretion, protein expression was induced for 4h with doxycycline before addition of 10 µg/ml Brefeldin A for 2 additional hours. Wah-out of cell-surface associated proteins was performed by incubation with 10 mg/ml Heparin for 5 min followed by 2 washes with PBS. To reduce PIP2 accessibility, cells were pre-treated with 10 mM neomycin. For cholesterol extraction experiments, cells were pre-treated for 30 minutes with 10 mM methyl-B-cyclodextrin. Plasma membrane targeting of 5phosphatase was performed by treatment with 100 nM rapamycin for 30 minutes.

### 5.3. Immunocytochemistry

Cells grown on coverslips were fixed with paraformaldehyde (4%, 10 min, room temperature) in PBS, permeabilized or not with Triton X-100 (0.3%, 5 min, room temperature) and saturated with PBS containing 10% FCS before incubation with primary antibodies (1 h, room temperature) and Alexa-labeled secondary antibodies (Thermo 1:500, 1 h, room temperature).

PIP2 staining was performed according to (Hammond et al., 2009). Cells grown on coverslips were washed in PBS and fixed (PBS, 4% PFA, 0.25% glutaraldehyde) for 15 minutes at RT and then on ice for 1h. All subsequent steps were performed on ice. After 3 washes (PBS, NH_4_Cl 50mM, 15 minutes each), saturation was proceeded in Buffer A (20mM Pipes, 137mM sodium chloride, 2.8mM potassium chloride. pH 6.8) supplemented with 0.5% Saponin and 5% FCS for 2h. Primary antibody (anti-PIP2, 2C11, Echelon, 5 µg/ml in Buffer A, 0.1% Saponin, 2% FCS) was added for 2h or overnight. After three washes (Buffer A, 0.1% Saponin), secondary antibody (AlexaFluor488 donkey anti mouse IgM 1/400, Life Technologies in Buffer A, 0.1% Saponin 2% FCS) was performed. Coadded for 1h. Coverslips were then washed three times (Buffer A, 0.1% Saponin) before post-fixation with PFA 2% in Buffer A (15 minutes on ice and then 10 minutes at RT). After rinsing with water, coverslips were dried and mounted.

### 5.4. Internalization

Cells (30,000 per well) were plated on μ-slide six-well plates (Ibidi). After 24 h, the medium was removed and cells were incubated with the fluorescent protein (1 *µ*M) diluted in DMEM for 30 minutes at 37°C before visualization. Cells were analyzed either directly to visualize the cell-surface staining or following addition of Trypan Blue (0.1% final concentration), an efficient quencher of all extracellular fluorescence and a marker of permeabilized cells, to visualize only the intracellular staining in live cells.

### 5.5. DNA constructs and recombinant proteins

All DNA constructs used in this study are listed in Table S2-3 and available upon request. His6-tagged recombinant proteins were produced in BL21 (DE3) grown in MagicMedia (Invitrogen) 24h, 28°C and purified on HisTrap columns (GE Healthcare) by Imidazole gradient elution on AKTA Prime according to manufacturer instructions. Following tag removal by incubation with PreScisson protease (6h, 4°C), the protein was purified on Heparin column (GE Healthcare), eluted by NaCl gradient elution and dialyzed for 2 days (20 mM phosphate buffer, 100 mM NaCl, pH 7.5). For protein labelling, 100 μM of dialyzed purified protein was incubated with a two folds molar excess of fluorescein isothiocyanate in carbonate buffer (50 mM pH 9.5, 100 mM NaCl) overnight at 4°C and free FITC was removed by dialysis (48 h, 4°C). The efficacy of FITC incorporation was controlled by SDS-PAGE and spectral analysis. The FITC:protein molecular ratio is controlled and ranges between 1.5 and 2 for all proteins.

### 5.6. Quantitative Secretion assay

Cells (13,000 per well) were plated on 96-well plates (Greiner Bio-one) coated with 1.5 µg/ml PolyDL-ornithine and induced for protein expression with doxycycline (1µg/ml). After 10 h, cells were transfected with the indicated plasmids and cultured for 24h. The medium was then replaced with fresh medium in the incubator and protein release was induced by biotin addition (100 µM final) and following Furmazin (Promega, 1/500) addition, luciferase activity was measured 1h later on a plate reader (Tristar, Berthold). Cells were then lysed in the presence of recombinant LgBiT (0.7 *µ*M final concentration) and Furimazin to measure intracellular protein expression. Rush-specific secretion was calculated by subtracting biotin-treated with biotin-untreated (control) wells for each condition and normalization with intracellular protein expression.

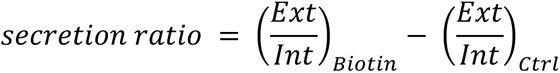

### 5.7. Quantitative Internalization assay

Cells (90,000 per well) were plated on 24-well culture dishes. After 24h, the protein was added to the medium and incubated for 30 minutes at 37°C. Following to washes with PBS, cells were collected by centrifugation after Trypsin treatment (0.25% final concentration) and resuspended in PBS 2% serum. Furimazin (1/500) was added to live cells and luciferase activity of the internalized protein was measured on a plate reader (Tristar, Berthold).

### 5.8. Imaging

Images of fixed / live cells were obtained with a CSU-W1 Yokogawa spinning disk coupled to a Zeiss Axio observer Z1 inverted microscope equipped with a sCMOS Hamamatsu camera and a 63× (1.4 Oil; WD:0.17mm) objective.

### 5.9. PWR

Self-assembled planar lipid bilayers used for PWR experiments were obtained from spontaneous fusion of a 3 mg/mL solution of small unilamellar liposomes (SUVs) composed of egg PC, egg PC/DOPS (3/1 mol/mol), egg PC/PIP_2_ (9/1 mol/mol) and egg PC/cholesterol/PIP_2_ (7/2/1 mol/mol/mol) into the silica surface of the PWR sensor (prism). PWR assays were performed by using a homemade PWR instrument (Harté et al., 2014). Light was generated from a polarized CW laser (He-Ne; wavelength of 632.8 nm) incident on the back surface of a thin metal film (Ag) deposited on a glass prism and coated with a layer of SiO2. The spectral angular resolution was *≤* 1 mdeg. PWR spectra corresponding to plots of reflected light intensity versus incident angle was excited with light whose electric vector was either parallel (s-polarized) or perpendicular (p-polarized) to the plane of the resonator surface. After formation and stabilization of the lipid bilayer (no further spectral changes with time), the protein was incrementally added to the PWR cell sample. Spectral changes were acquired for both polarizations. The system was let to equilibrate before each peptide addition. PWR being sensitive to the optical properties of material deposited on the resonator surface (so peptide bound to the lipid bilayer), interference from the material present in the bulk solution (non-bound) is unlikely. Apparent dissociation constants (KD) were obtained by plotting the resonance minimum position as a function of the peptide concentration and by fitting the plot through a hyperbolic binding function using GraphPad Prism™ version 5.0a (GraphPad Software, San Diego, California, US).

### 5.10. Cell Fractionation

Cells were lysed by Dounce homogenization in hypotonic lysis buffer (Hepes 10mM, KCl 10mM, MgCl_2_ 3 mM, EGTA 1mM, Protease inhibitors) subsequently adjusted to 0.25M Sucrose. Following nuclei removal (1,000 *g*, 10 min), membranes were pelleted by centrifugation (150,000 *g*, 30 min) and washed with sucrose lysis buffer before being pelleted by a second round of centrifugation. Membrane pellet was resuspended in Sucrose Lysis buffer and treated or not with freshly prepared neomycin (10 mM final concentration) 30 min at 4°C. Membrane samples were diluted 10 folds in sucrose lysis buffer and released and membrane-associated proteins were separated by centrifugation (150,000 *g*, 30 min). Pellets were resuspended directly in Laemmli buffer and supernatant proteins were precipitated (0.02% Deoxycholate, 4% TCA, 30 min à 4°C) before resuspension in Laemmli buffer in order to load equivalent fractions of each sample on SDS PAGE.

### 5.11 Statistical Analysis

Data were analyzed with GraphPad Prism 6 and expressed as the mean +/- SEM. Two groups were compared with student’s t-test. For multiple group comparison, one-way analysis of variance (ANOVA) followed by Tukey’s post-tests was performed. P-values < 0.05 were considered to be statistically significant.

* p< 0.05; ** p < 0.01; *** p < 0.001; **** p < 0.0001

## Acknowledgments

We thank S.Taylor for providing us with the FLpIn-TREX HeLa cell line. and A. Echard for providing with plasmids and S. Vriz and M. Volovitch for fruitful discussions and careful rereading of the manuscript:

## Competing interests

The authors declare no competing interests.

## Funding

This work supported by CNRS, INSERM, Collège de France, Université de Paris, Labex PSL Memolife and Agence Nationale de la Recherche, (ANR-17-CE11-0050).

---

**Table S1:**
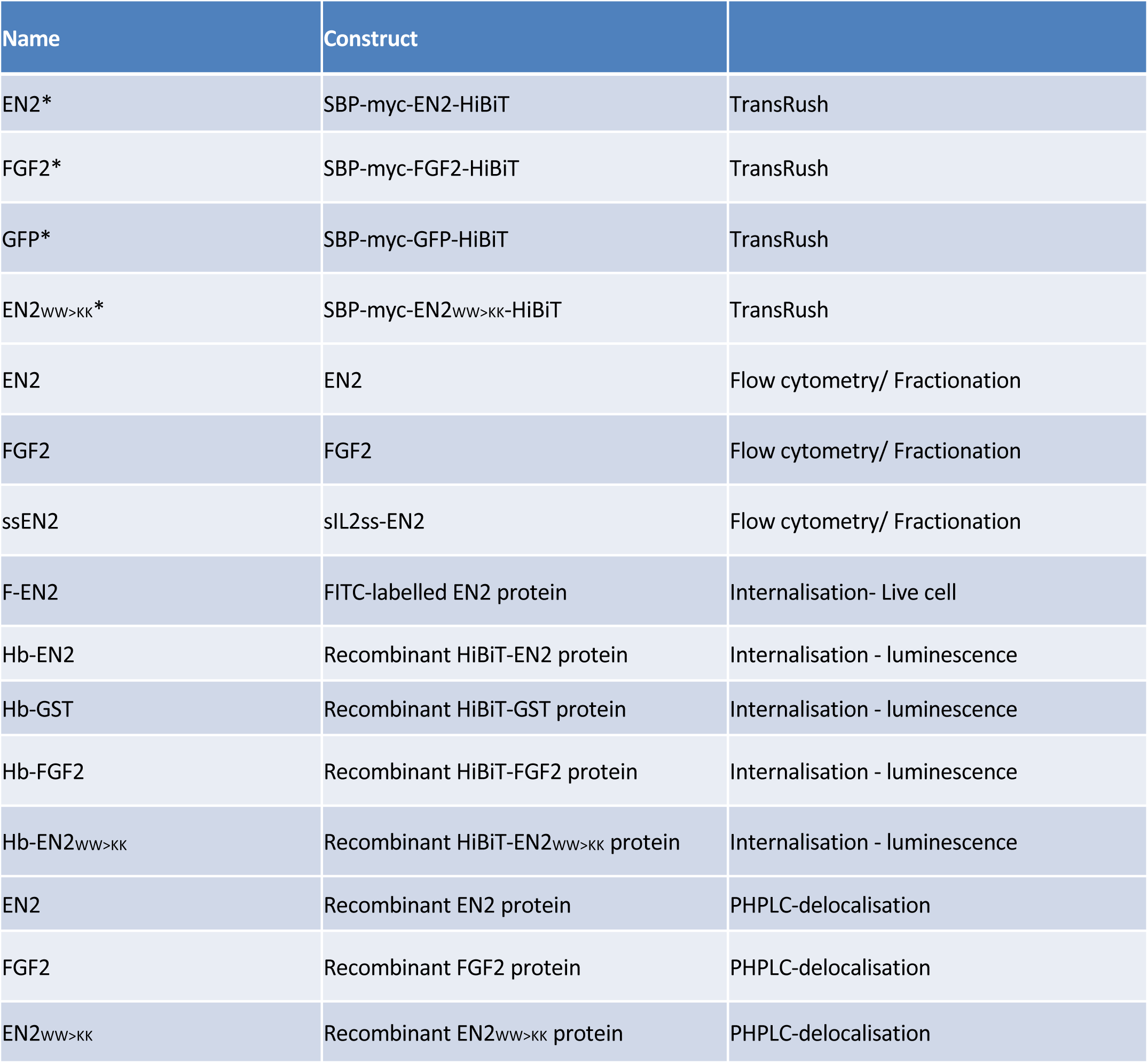
Abbreviations of protein names used in the study

| Name | Construct |  |
| --- | --- | --- |
| EN2* | SBP-myc-EN2-HiBiT | TransRush |
| FGF2* | SBP-myc-FGF2-HiBiT | TransRush |
| GFP* | SBP-myc-GFP-HiBiT | TransRush |
| EN2<sub>WW>KK</sub>* | SBP-myc-EN2<sub>WW>KK</sub>-HiBiT | TransRush |
| EN2 | EN2 | Flow cytometry/ Fractionation |
| FGF2 | FGF2 | Flow cytometry/ Fractionation |
| ssEN2 | sIL2ss-EN2 | Flow cytometry/ Fractionation |
| F-EN2 | FITC-labelled EN2 protein | Internalisation- Live cell |
| Hb-EN2 | Recombinant HiBiT-EN2 protein | Internalisation - luminescence |
| Hb-GST | Recombinant HiBiT-GST protein | Internalisation - luminescence |
| Hb-FGF2 | Recombinant HiBiT-FGF2 protein | Internalisation - luminescence |
| Hb-EN2<sub>WW>KK</sub> | Recombinant HiBiT-EN2<sub>WW>KK</sub> protein | Internalisation - luminescence |
| EN2 | Recombinant EN2 protein | PHPLC-delocalisation |
| FGF2 | Recombinant FGF2 protein | PHPLC-delocalisation |
| EN2<sub>WW>KK</sub> | Recombinant EN2<sub>WW>KK</sub> protein | PHPLC-delocalisation |

**Table S2:**
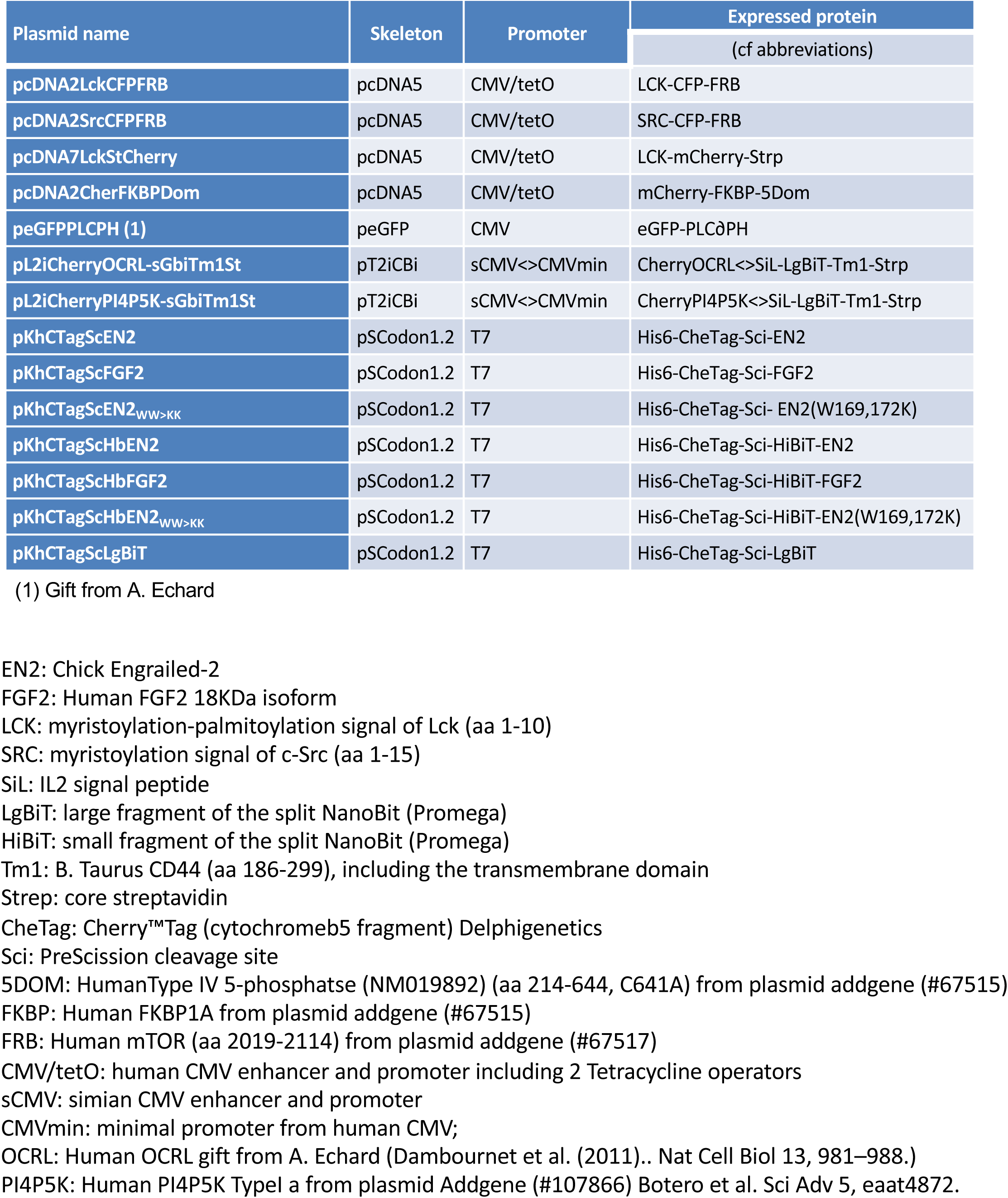
DNA constructs

**Table S3:** Plasmid used for the generation of HeLa Cell clones

| Plasmid name | Skeleton | Promoter | Expressed Protein<br>(cf abbreviation) |
| --- | --- | --- | --- |
| pcDNA7EN2 | pcDNA5 | CMV/tetO | EN2 |
| pcDNA7ssEN2 | pcDNA5 | CMV/tetO | Sil-EN2 |
| pcDNA7MycFGF2 | pcDNA5 | CMV/tetO | Myc-FGF2 |
| pcDNA7EN2* | pcDNA5 | CMV/tetO | SBP-myc-EN2-HiBiT |
| pcDNA7FGF2* | pcDNA5 | CMV/tetO | SBP-myc-FGF2-HiBiT |
| pcDNA7GFP* | pcDNA5 | CMV/tetO | SBP-myc-GFP-HiBiT |
| pcDNA7EN2<sub>WW>KK</sub>* | pcDNA5 | CMV/tetO | SBP-myc-EN2(W169,172K)-HiBiT |
| PcDNA7LgBiT | pcDNA5 | CMV/tetO | LgBiT |

EN2: Chick Engrailed-2
FGF2: Human FGF2 18KDa isoform
Sil: IL2 signal peptide
LgBiT: large fragment of the split NanoBit (Promega)
HiBiT: small fragment of the split NanoBit (Promega)
Strep: core streptavidin
SBP: streptavidin-binding protein
myc: 10-aa myc tag
CMV/tetO: human CMV enhancer and promoter including 2 Tetracycline operators

**Table S4.** Statistical data of PWR experiments

**Fig. 1E**
|  | PC | PC/PS | PC/PIP<sub>2</sub> |
| --- | --- | --- | --- |
| **Mean** | **6.333** | **2.767** | **0.0750** |
| **Std. Deviation** | **1.201** | **0.7506** | **0.01323** |
| Std. Error | 0.6936 | 0.4333 | 0.007638 |

**Fig 5A**
|  | PC/PIP<sub>2</sub> | PC/PIP<sub>2</sub>/Chol |
| --- | --- | --- |
| **Mean** | **0.0750** | **0.0110** |
| **Std. Deviation** | **0.01323** | **0.003606** |
| Std. Error | 0.007638 | 0.002082 |

**Fig. 7E**
|  | PC | PC/PIP<sub>2</sub> |
| --- | --- | --- |
| **Mean** | **20.33** | **2.533** |
| **Std. Deviation** | **4.509** | **0.6506** |
| Std. Error | 2.603 | 0.3756 |

**Fig. S1B**
|  | PC | PC/PS | PC/PIP<sub>2</sub> |
| --- | --- | --- | --- |
| **Mean** | **49.33** | **41.00** | **0.1200** |
| **Std. Deviation** | **7.024** | **5.568** | **0.02646** |
| Std. Error | 4.055 | 3.215 | 0.01528 |

**Fig S1:**
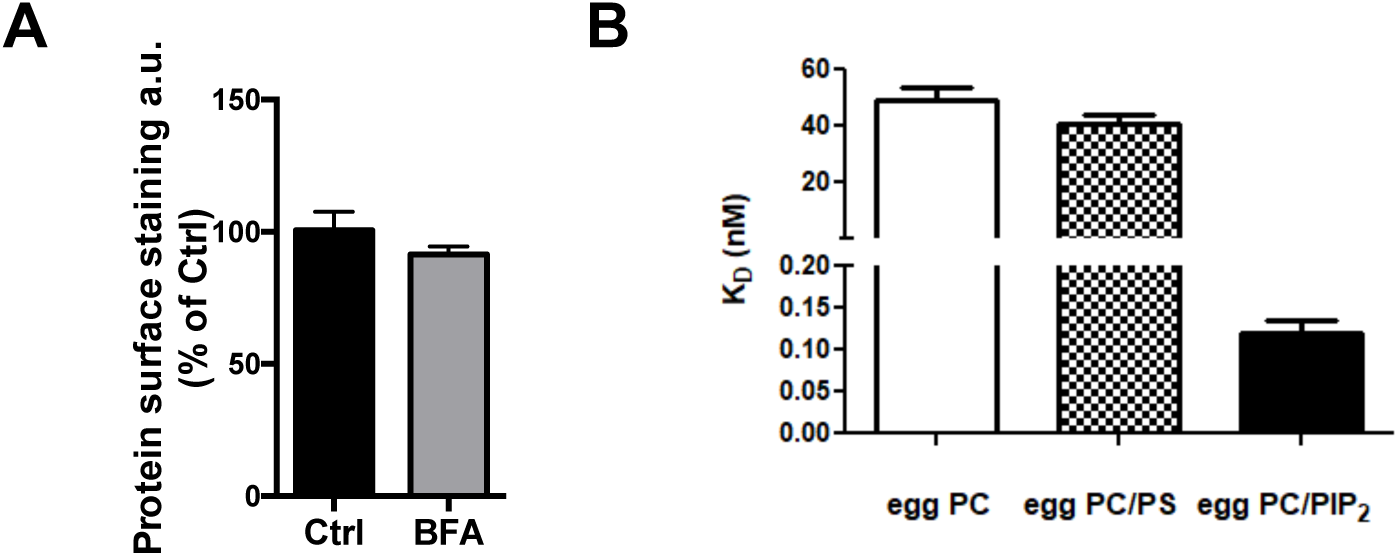
Behaviour of FGF2 in secretion and PIP2 Interaction assays. (A) Insensitivity of FGF2 secretion to Brefeldin A treatment as in Fig. 1A. (B) Binding affinity of FGF for lipid membranes composed of egg PC, egg PC/DOPS (3/1 mol/mol) and egg PC/PIP2 (9/1 mol/mol) as in FIG. 1E.

**Fig S2:**
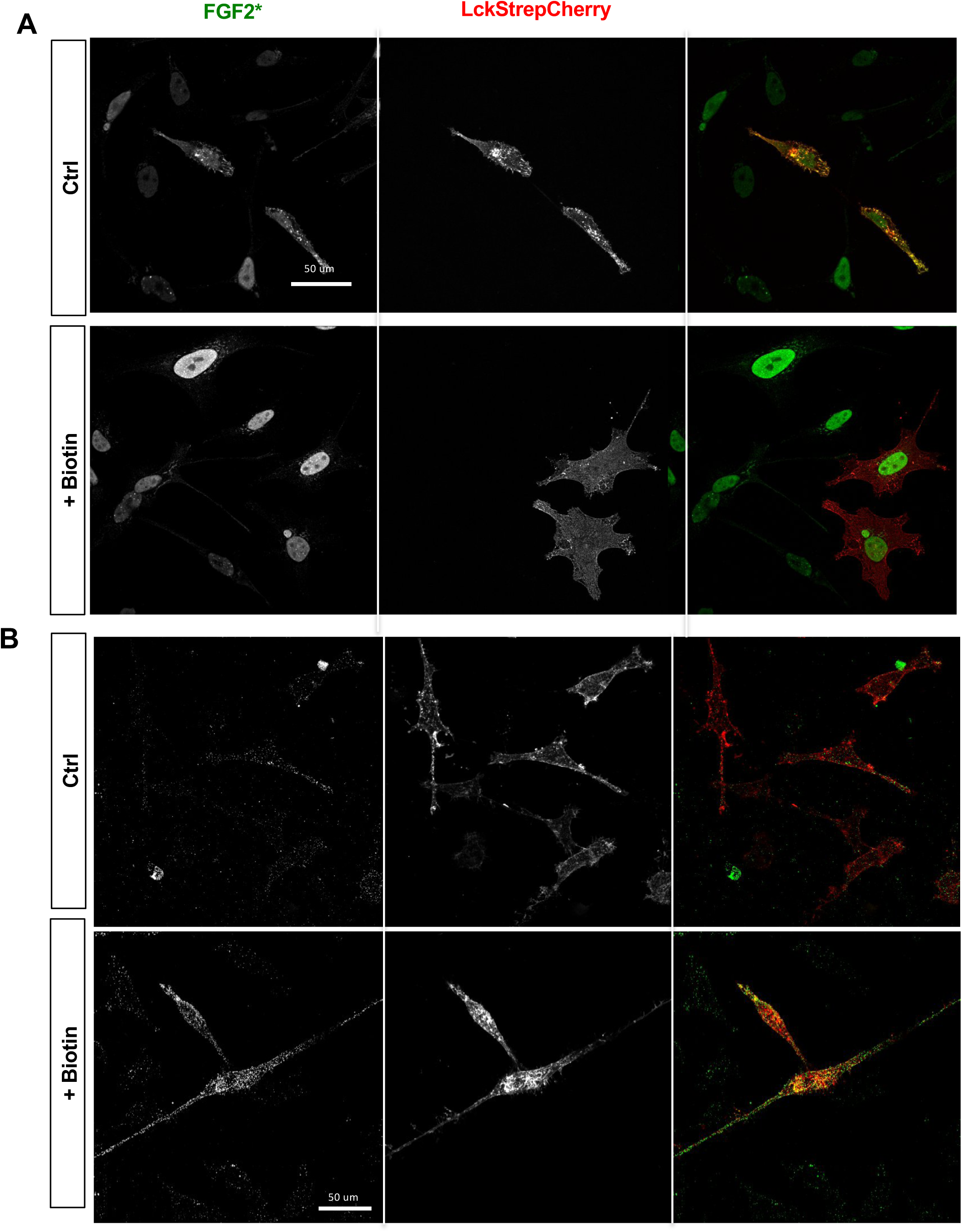
Validation of Rush assay for FGF2 in permeabilized and unpermeabilized cells. Stable HeLa cell lines expressing FGF2* were transfected with a LCKStrepCherry construct and treated or not with Biotin (60 min, 100 µM). A) Intracellular and B) extracellular FGF2* were detected by immunofluorescence using permeabilised or unpermeabilized conditions respectively.

**Fig S3:**
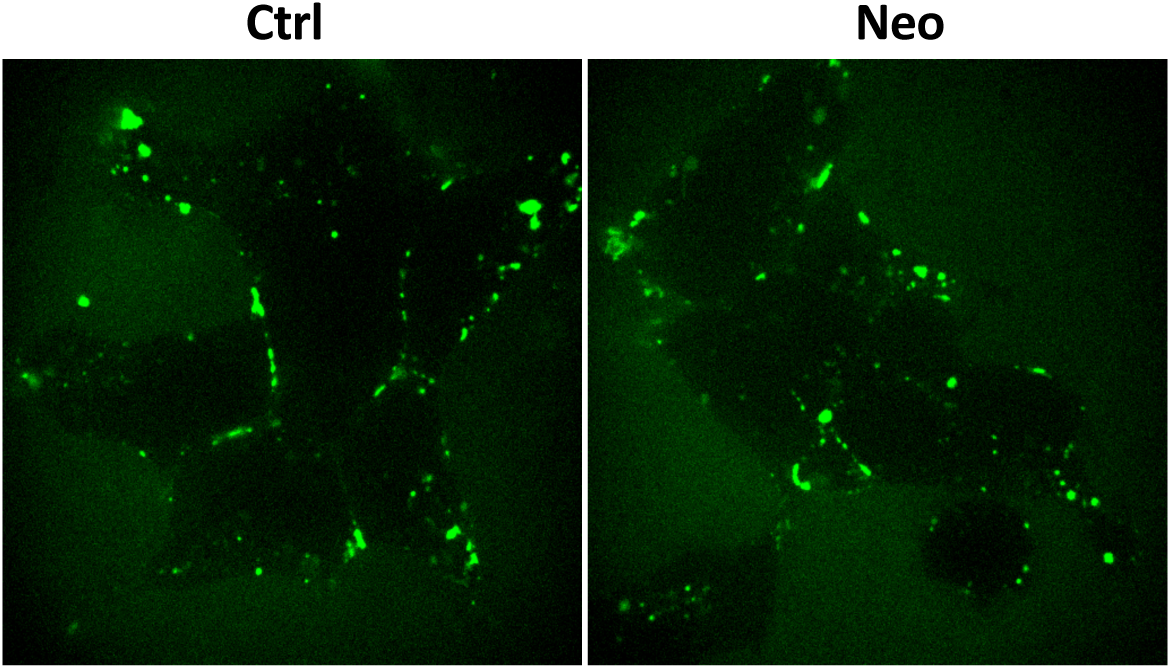
Neomycin treatment does not affect EN2 extracellular localisation. Cells pretreated with neomycin (neo) or not (ctrl) and incubated with F-EN2. Extracellular F-EN2 was visualised (before trypan blue addition).

**Fig S4:**
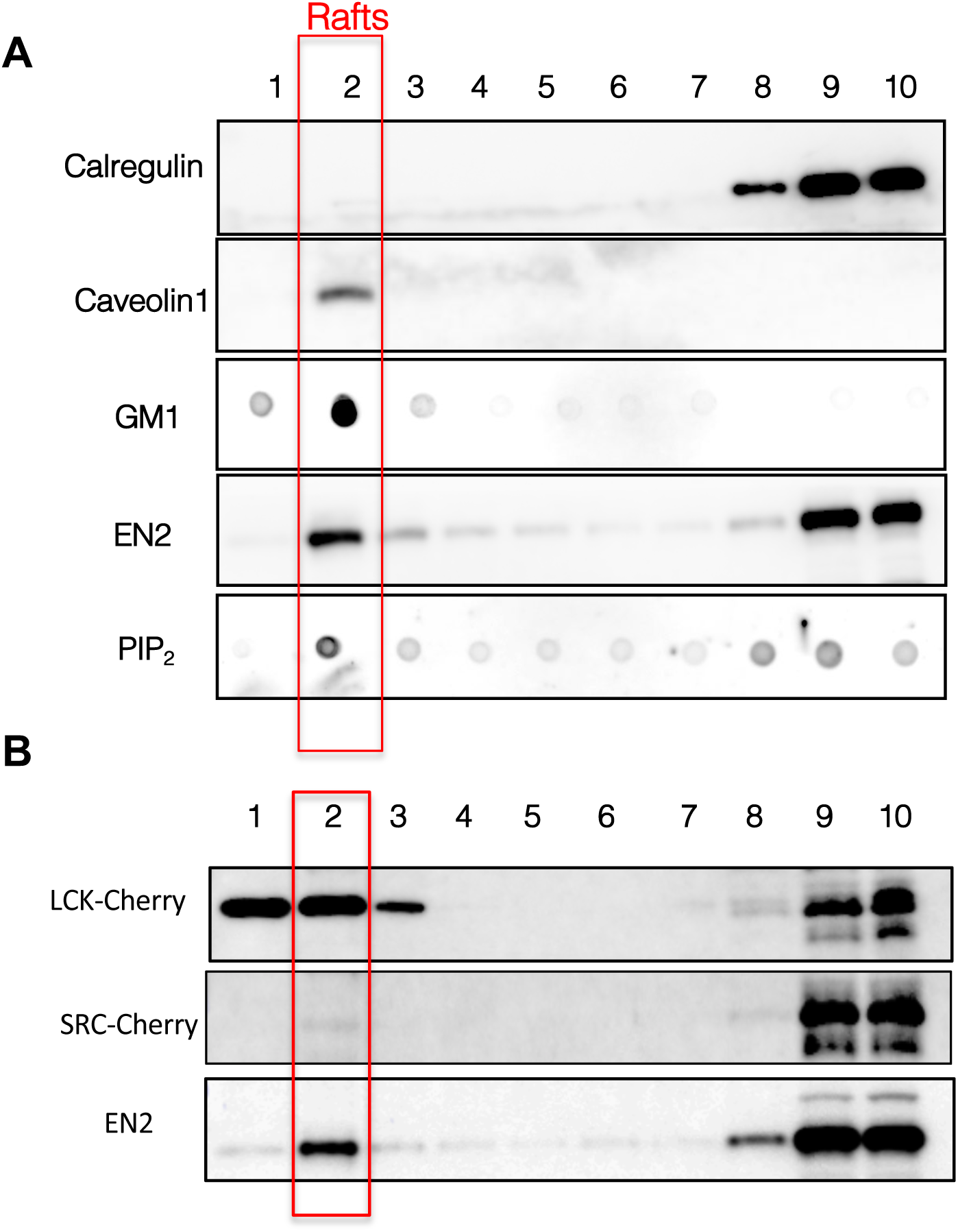
Role of raft associated PIP2. (A) The raft (cholesterol-enriched) membrane fraction purified from EN2 cell line, characterized by the presence of Caveolin1 and GM1 ganglioside (Cholera toxin staining) and the absence of ER marker Calregulin, is enriched both in PIP_2_ (KT10 antibody staining) and EN2. (B) The LCK anchor, but not the SRC one, drive a mCherry reported toward raft fractions Raft separation was performed as described in **Yamaguchi, H., Yamaguchi, H., Shiraishi, M., Shiraishi, M., Fukami, K., Tanabe, A., Ikeda-Matsuo, Y., Ikeda Matsuo, Y., Naito, Y., Naito, Y., et al.** (2009). MARCKS regulates lamellipodia formation induced by IGF-I via association with PIP2 and beta-actin at membrane microdomains. *J Cell Physiol* **220**, 748–755. Briefly, Cells extracts were resuspened in (Hepes 10mM, KCl 10mM, MgCl2 3 mM, EGTA 1mM) +I.P.+ 0,25M saccharose+1%Brij-35) incubated on ice for 15 min and sonicated. The Post nuclear supernatant adjusted to 35% optiprep was loaded at the bottom of a 5-30 % Optiprep gradient and centrifuged (250 000*g*, 2h). 10 fractions (1-Light, 10-Heavy) were collected and processed for western blot analysis.

**Fig S5.**
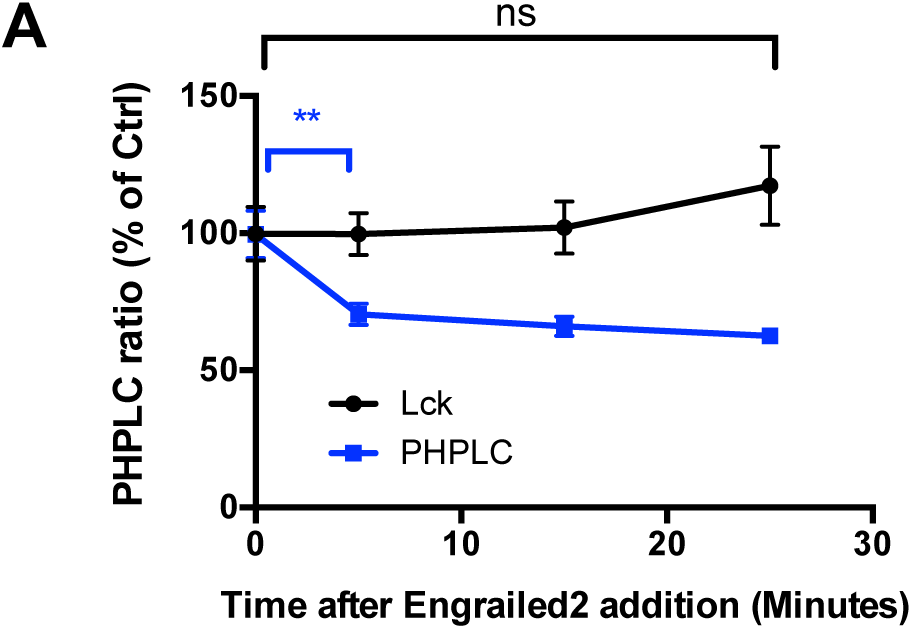
Differential sensitivity of PHPLC and LCK anchor delocalisation to EN2 addition. Quantification of PHPLC or LCK-Cherry delocalisation from the plasma membrane following extracellular addition of 2 µM EN2.

